# Zero-Shot Design of a Biobetter Cetuximab: Enhanced EGFR Affinity with Preserved Developability

**DOI:** 10.64898/2026.05.05.722890

**Authors:** Iddo Weiner

## Abstract

Cetuximab is a chimeric IgG1 monoclonal antibody that has been a cornerstone therapy for EGFR-driven malignancies for nearly two decades. Its therapeutic activity is governed by competitive displacement of endogenous EGFR ligands, making binding affinity a direct determinant of clinical efficacy. We applied ConvergeAB™, a target-aware antibody design platform, in a fully zero-shot configuration to generate a biobetter version of cetuximab. The lead Converge-designed antibody binds EGFR with a mean KD of 315 pM — approximately 2.1-fold tighter than cetuximab (673 pM) and 4.4-fold tighter than a recently published, computationally designed anti-EGFR antibody from Cradle Bio (1.38 nM). The affinity gain arises from six substitutions that leave the global paratope architecture intact (Cα RMSD 0.15 Å vs cetuximab) and instead optimize the binding interface through localized packing and electrostatic adjustments. A panel of biophysical and developability assays — HIC, DLS, DSF, and PSR ELISA — shows that the Converge variant matches or exceeds cetuximab on monomericity, monodispersity, polyspecificity, and thermal stability, while remaining within a developable hydrophobicity envelope. Together, these data demonstrate that a single zero-shot ConvergeAB™ campaign can deliver a biobetter molecule with significantly improved affinity and a clean developability profile, without compromising the parental antibody’s drug-like properties.

## Introduction

Epidermal Growth Factor Receptor (EGFR) is a transmembrane tyrosine kinase that plays a central role in regulating cell proliferation, survival, and differentiation. Upon binding of its natural ligands, such as epidermal growth factor (EGF), EGFR undergoes dimerization and autophosphorylation, triggering downstream signaling cascades. Dysregulation of EGFR — through overexpression or mutation — is a well-established driver of oncogenesis. Elevated EGFR activity has been implicated across multiple solid tumors, most notably in colorectal cancer, head and neck squamous cell carcinoma (HNSCC), and non-small cell lung cancer (NSCLC). In these contexts, aberrant EGFR signaling promotes uncontrolled cell growth, resistance to apoptosis, and enhanced metastatic potential, making it a compelling therapeutic target.^1^

Cetuximab is a chimeric monoclonal antibody designed to target the extracellular domain of EGFR. By binding to the receptor with high specificity, cetuximab sterically blocks ligand interaction, thereby preventing receptor activation and downstream signaling. Clinically, cetuximab has become a cornerstone therapy in EGFR-driven malignancies, particularly in metastatic colorectal cancer and recurrent or metastatic HNSCC and is widely used in combination with chemotherapy or radiation, with global adoption supported by regulatory approvals.^2^

At the molecular level, the therapeutic efficacy of cetuximab is fundamentally governed by its competition with endogenous ligands such as EGF for binding to EGFR. This competitive interaction is dictated by binding affinity, commonly quantified by the equilibrium dissociation constant (K_D_). A lower K_D_ reflects tighter binding and greater competitive advantage. In tumors where ligand concentrations are high, cetuximab must effectively outcompete EGF to suppress receptor activation; insufficient affinity allows residual signaling and contributes to suboptimal response and disease progression. Optimizing K_D_ is therefore not merely a biochemical consideration but a critical determinant of clinical efficacy, influencing both depth and durability of response in patients.^3^

In this work, we apply ConvergeAB™, a target-aware antibody design platform, in a fully zero-shot configuration to generate a biobetter version of cetuximab. Throughout this work, we benchmark our design against two reference molecules. The first is cetuximab itself — the clinical standard of care for anti-EGFR antibody therapy and the seed of our affinity-maturation campaign. The second is a recently published anti-EGFR antibody from Cradle Bio, which won a public protein-design competition organized by Adaptyv Biosystems and represents a contemporary, computationally designed comparator.^4^ Including both references allows us to position the Converge-designed molecule against the in-market gold standard and against the current frontier of de novo computational antibody design.

## Results

### Zero-shot affinity maturation with ConvergeAB™

To generate a biobetter version of cetuximab, we prompted ConvergeAB™ with the original cetuximab sequence together with the EGFR target sequence. No task-specific or ad-hoc training was performed prior to the initiation of this affinity-maturation campaign, making it a fully zero-shot effort. ConvergeAB™ is a multi-step platform in which sequence diversity is introduced into a seed antibody using a proprietary protein language model, generating ∼100,000 target-aware candidates. These candidates are subsequently filtered through a series of explicit and orthogonal predictors that evaluate key properties — including binding affinity, structural compatibility (docking), thermal stability, and solubility. The top 10 predicted candidates from this campaign were selected for experimental validation. Each variant was expressed and screened for binding by single-concentration surface plasmon resonance (SPR), and the highest-performing candidate was advanced for full kinetic characterization.

### The Converge-designed antibody binds EGFR with significantly higher affinity than cetuximab and Cradle

All three antibodies — Converge, cetuximab, and Cradle — were produced as full-length human IgG1 (K214R; consistent with the clinical form of cetuximab). For rigorous head-to-head comparison, the three antibodies were analyzed side-by-side in two independent multi-cycle kinetic SPR experiments, each using six analyte concentrations of recombinant human EGFR-His. Across all measured parameters (k_on_, k_off_, and K_D_), and in both repeats, the Converge-designed antibody consistently outperformed both reference molecules (Table 1; Figure 2).

**Table 1.** SPR kinetic parameters for the three anti-EGFR antibodies. Each repeat represents an independent, parallel measurement of all three antibodies under identical conditions. Lower KD indicates tighter binding.

| Antibody | Repeat | $k_{on}$ (1/Ms) | $k_{off}$ (1/s) | $K_D$ (pM) |
| --- | --- | --- | --- | --- |
| Converge | #1 | $5.76 \times 10^6$ | $1.28 \times 10^{-3}$ | 222 |
| Converge | #2 | $3.10 \times 10^6$ | $1.27 \times 10^{-3}$ | 409 |
| Converge | Mean | $4.43 \times 10^6$ | $1.275 \times 10^{-3}$ | **315** |
| Cetuximab | #1 | $3.23 \times 10^6$ | $1.81 \times 10^{-3}$ | 560 |
| Cetuximab | #2 | $2.25 \times 10^6$ | $1.77 \times 10^{-3}$ | 786 |
| Cetuximab | Mean | $2.74 \times 10^6$ | $1.79 \times 10^{-3}$ | **673** |
| Cradle | #1 | $3.92 \times 10^6$ | $4.32 \times 10^{-3}$ | 1,100 |
| Cradle | #2 | $2.53 \times 10^6$ | $4.21 \times 10^{-3}$ | 1,660 |
| Cradle | Mean | $3.22 \times 10^6$ | $4.265 \times 10^{-3}$ | **1,380** |

**Figure 1.**
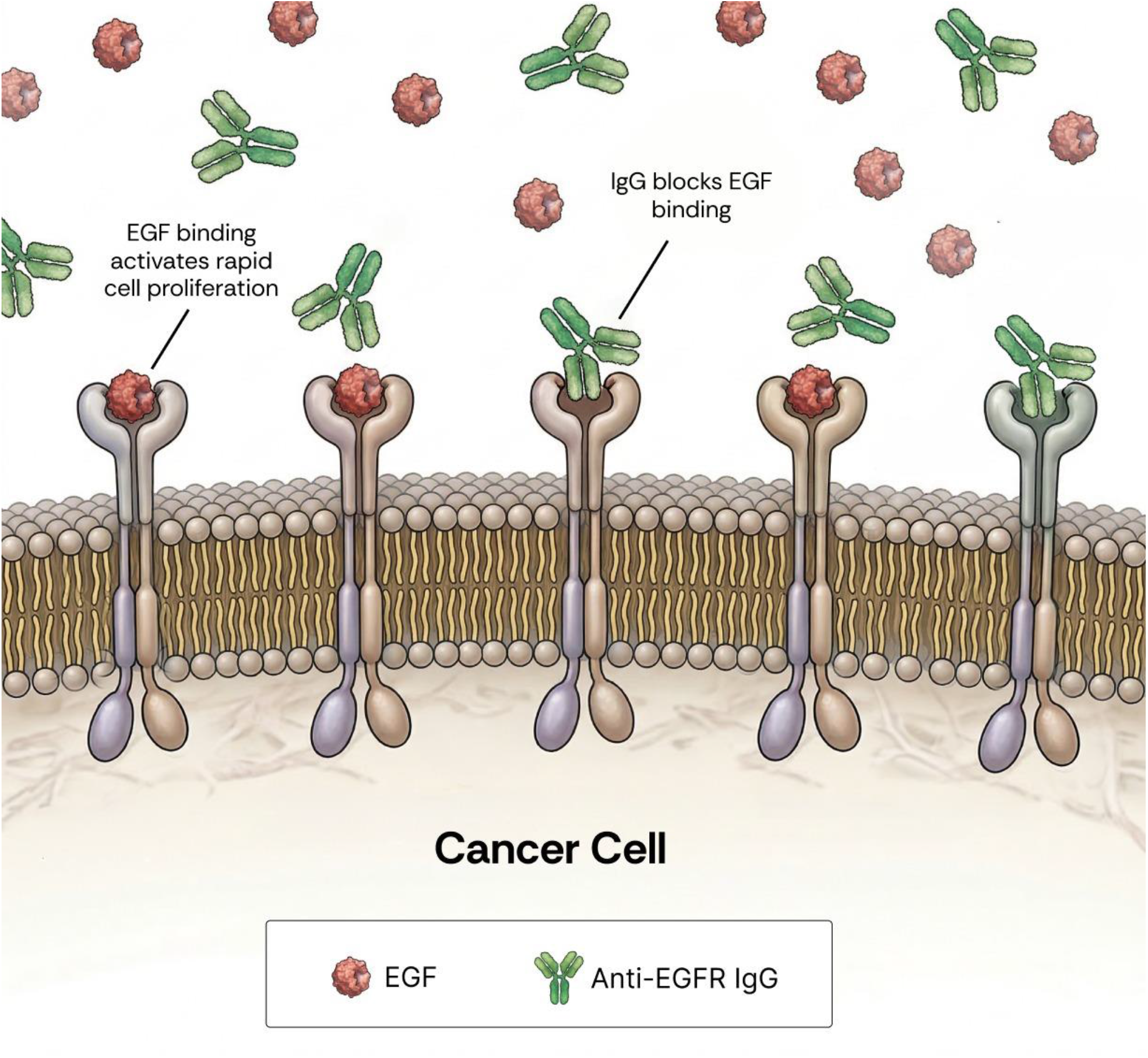
Competitive binding at EGFR is a key determinant of the therapeutic efficacy of anti-EGFR antibodies. Epidermal growth factor (EGF, red) and anti-EGFR monoclonal IgG (green) compete for binding to the EGFR transmembrane receptor on the surface of cancer cells. Engagement of EGFR by EGF activates proliferative signaling pathways that drive tumor growth and progression. Antibody binding blocks ligand-induced receptor activation, leading to growth inhibition, cell-cycle arrest, or apoptosis. The outcome of this competition is governed by binding kinetics, with lower-KD antibodies exhibiting superior competitive advantage over endogenous ligands.

**Figure 2.**
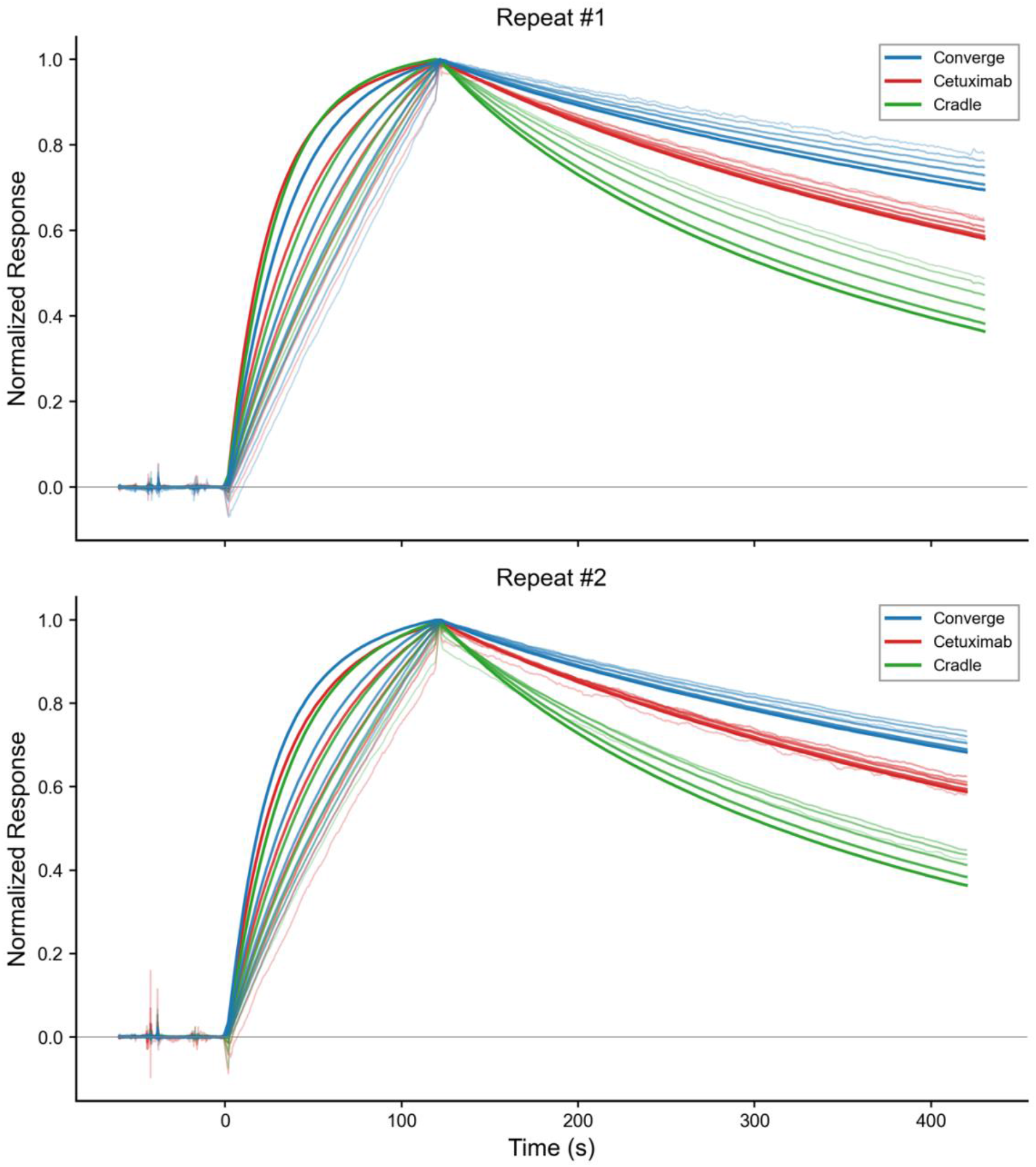
SPR sensorgrams from both repeats. Curves are Y-normalized to enable side-by-side comparison across runs. EGFR was used as the analyte at the following concentrations (nM): 0.20, 0.39, 0.78, 1.6, 3.1, 6.2 (and 12.5 in the broader concentration set). Curve thickness corresponds to analyte concentration, with the boldest curve representing the highest concentration. The slower dissociation phase of Converge (blue) relative to cetuximab (red) and Cradle (green) is evident in both repeats.

The Converge antibody achieved a mean KD of 315 pM — ∼2.1-fold tighter than cetuximab (mean 673 pM) and ∼4.4-fold tighter than Cradle (mean 1.38 nM). The improvement was driven by both faster association (1.6x higher k_on_ than cetuximab) and slower dissociation (1.4x lower k_off_). Notably, Cradle’s k_off_ was ∼3.4-fold faster than Converge’s, indicating substantially shorter target residence and explaining its weaker overall K_D_ despite a competitive k_on_.

### Six edits drive the affinity gain without altering global architecture

Sequence analysis revealed that the Converge antibody differs from cetuximab at six positions, comprising four substitutions to alanine (VH T61A, VH S87A, VL V9A, VL N93A) and two asparagine-to-aspartate substitutions (VH N88D, VL N32D), distributed across both framework and CDR regions (Figure 3). Pairwise sequence identity between Converge and cetuximab is 97.5% in VH and 97.2% in VL. For comparison, Cradle is also closely related to cetuximab at the sequence level (VH 95.0% identity, VL 96.3%), with edits concentrated in framework regions alone (VH positions 5, 70, 71, 73, 87, 88 and VL positions 45, 49, 79, 80).

**Figure 3.**
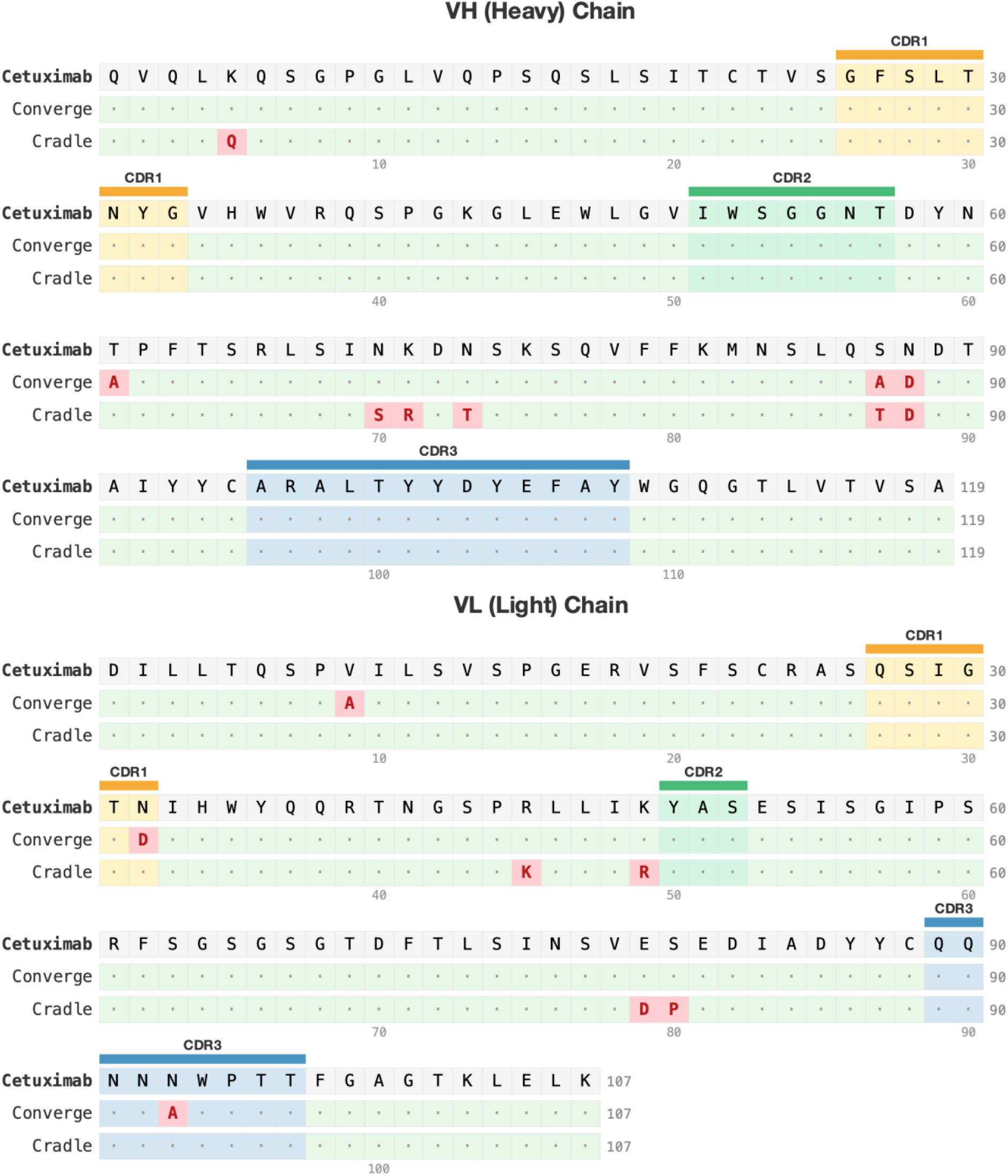
Sequence alignment of cetuximab, the Converge-designed anti-EGFR antibody, and Cradle’s anti-EGFR antibody. CDR1, CDR2, and CDR3 regions (IMGT definitions) are annotated. Mutated positions are highlighted; the Converge variant carries six edits relative to cetuximab (VH T61A, VH S87A, VH N88D, VL V9A, VL N32D, VL N93A) distributed across framework and CDR regions.

To assess whether these affinity gains were associated with large-scale structural remodeling, we predicted the 3D structures of the Converge variant, cetuximab, and Cradle and aligned them. The Converge–cetuximab pairwise Cα RMSD was 0.15 Å (VH 0.14 Å, VL 0.16 Å), indicating that the Converge variant preserves the global cetuximab binding architecture (Figure 4). Cradle was likewise structurally close to cetuximab (RMSD 0.17 Å). This is an important observation, as it argues against a wholesale change in fold or binding mode and instead supports a mechanism in which affinity improvement is driven by highly localized structural and energetic optimization within an otherwise conserved paratope.

**Figure 4.**
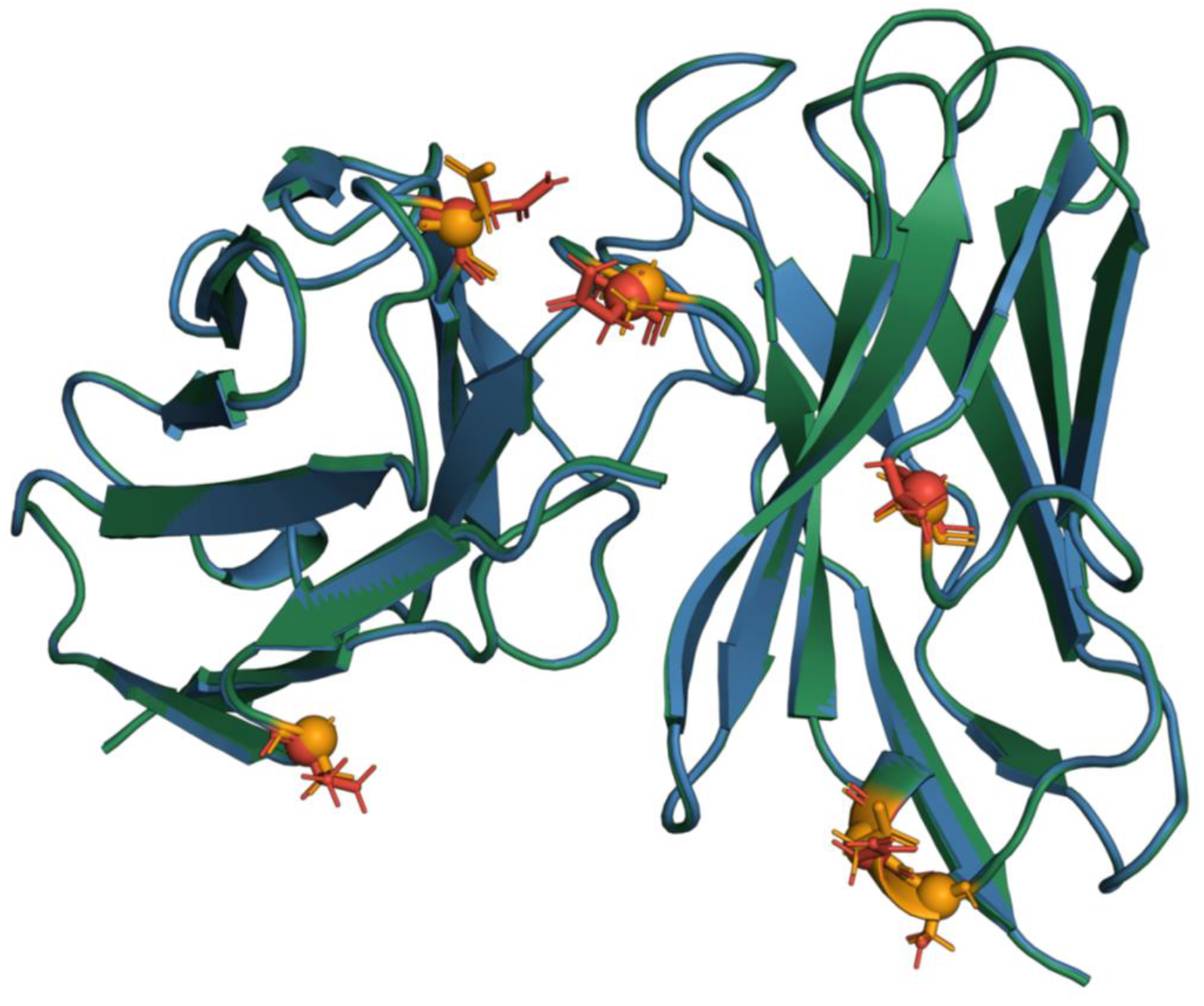
Structural superposition of cetuximab and the Converge-designed anti-EGFR antibody. Mutation sites relative to cetuximab are highlighted as red sticks. Despite the introduced edits, the global Fv architecture is highly conserved (pairwise Cα RMSD = 0.15 Å, VH Cα RMSD = 0.14 Å, VL Cα RMSD = 0.16 Å).

This interpretation is consistent with the nature of the Converge edits. The alanine substitutions are expected to reduce side-chain bulk and remove potentially suboptimal polar functionality, which can improve local packing, decrease conformational heterogeneity, and favor a more pre-organized binding-competent state. Likewise, the asparagine-to-aspartate substitutions introduce negative charge and altered hydrogen-bonding geometry, which may strengthen electrostatic complementarity with EGFR or stabilize productive local conformations. The structural comparison further supports this localized mechanism: although the global RMSD remains minimal, the two CDR edits appear to act through different routes. The N32D substitution in CDR-L1 represents a semi-conservative polar-for-polar exchange at a solvent-exposed position, where altered charge and hydrogen-bond geometry can refine electrostatic complementarity with EGFR without requiring backbone rearrangement. CDR-L3, by contrast, accounts for the bulk of the resolvable structural change: the VL N93A substitution produces a 0.66 Å shift at the Cα position and displaces neighboring Trp94 by 0.5 Å, and the marginally larger VL Cα RMSD (0.16 Å versus 0.14 Å for VH) tracks this loop-apex repacking. These are small but meaningful local rearrangements, especially in an antibody paratope, where sub-angstrom movements can measurably alter side-chain presentation, loop packing, and epitope complementarity without changing the overall fold.

Taken together, the sequence and structure analyses support a coherent model for the experimentally confirmed affinity improvement. Rather than arising from major structural divergence from cetuximab, the Converge sequence retains the parental architecture while introducing a set of cooperative micro-adjustments that fine-tune the binding interface — consistent with an affinity-maturation mechanism based on paratope preorganization, improved local packing, and targeted electrostatic optimization.

### Developability profile

Affinity gains are clinically valuable only when accompanied by acceptable biophysical behavior. To assess whether the Converge edits preserved cetuximab’s drug-like properties, we evaluated all three antibodies on an orthogonal panel of developability assays: hydrophobic interaction chromatography (HIC) for surface hydrophobicity, dynamic light scattering (DLS) for particle size and dispersity, differential scanning fluorimetry (DSF) for thermal stability, and a polyspecificity reagent (PSR) ELISA panel for non-specific binding. All three antibodies were produced from the same expression platform, and assays were run side-by-side with internal benchmark controls for each readout. Across all four assays, the Converge antibody matched or outperformed cetuximab and was generally cleaner than Cradle.

#### Hydrophobicity (HIC)

Surface hydrophobicity was measured by HIC-HPLC on a Thermo MabPac HIC-Butyl column (Table 2). Trastuzumab (anti-HER2 IgG1) was included in every run as the relative-retention-time (RRT = RTsample / RTtrastuzumab) reference; trastuzumab is a clinically approved IgG1 with a favorable well-characterized HIC profile.

**Table 2.** HIC retention times for the three antibodies. RT, retention time at the main 214 nm peak; RRT, relative retention time vs. trastuzumab control (RT = 6.473 min). Higher RRT indicates greater surface hydrophobicity. Main-peak Area% (214 nm) reports purity / monomer-dominance of the chromatogram.

| Antibody | RT (min) | RRT (vs trastuzumab) | Main peak Area% |
| --- | --- | --- | --- |
| Converge | 8.108 | 1.253 | 98.21 |
| Cetuximab | 6.967 | 1.076 | 95.94 |
| Cradle | 8.780 | 1.356 | 99.23 |
| Trastuzumab (control) | 6.473 | 1.000 | — |

All three antibodies eluted as predominantly single, sharp peaks (>95% main-peak area at 214 nm), with no evidence of aggregation- or fragmentation-related satellite peaks at developability-relevant levels. The Converge variant exhibits modestly higher hydrophobicity than cetuximab (RRT 1.25 vs 1.08), consistent with the four hydrophobic alanine substitutions introduced relative to cetuximab. Cradle is the most hydrophobic of the three (RRT 1.36). All values fall within the developable range typical for clinical-stage IgG1s; cetuximab itself is a marketed drug, providing the relevant in-class reference.

#### Hydrodynamic size and dispersity (DLS)

DLS was performed on a Wyatt DynaPro Plate Reader at 25 °C in water, with two replicate wells per antibody plus a trastuzumab control (Table 3). Trastuzumab serves as the same role here as in HIC: a well-behaved, clinically approved IgG1 monomer that anchors the expected radius and dispersity of an “uneventful” intact antibody on this instrument. A test antibody whose DLS readouts approach trastuzumab’s is expected to be similarly monomeric and monodisperse in solution; substantial deviation in radius or %PD signals oligomerization, aggregation, or inhomogeneity.

**Table 3.** DLS results (mean of two replicates). Lower hydrodynamic radius and lower %PD reflect a more compact, monodisperse population. Normalized intensity is reported for completeness but is sensitive to concentration and is not a developability metric on its own.

| Antibody | Radius (nm) | Diameter (nm) | %PD | Normalized Intensity (Cnt/s) |
| --- | --- | --- | --- | --- |
| Converge | 5.85 | 11.65 | 3.95 | $1.12 \times 10^7$ |
| Cetuximab | 6.80 | 13.55 | 10.45 | $1.92 \times 10^7$ |
| Cradle | 7.40 | 14.75 | 10.05 | $4.52 \times 10^7$ |
| Trastuzumab (control) | 5.70 | 11.35 | 5.45 | $1.32 \times 10^7$ |

The Converge variant shows the smallest hydrodynamic radius (5.85 nm) and the lowest polydispersity (3.95% PD) of the three test antibodies — values that are within the trastuzumab-control envelope and indicate a clean, monomeric, monodisperse population. Cetuximab and Cradle both run at ∼6.8–7.4 nm radius with ∼10% PD, consistent with intact IgG monomer plus a small number of higher-order species or polydispersity. The Cradle replicate-1 normalized intensity (7.2 × 10^7^ Cnt/s) is an outlier vs. its replicate 2 (1.8 × 10^7^), suggesting a low-level aggregate spike in one well — a possible early developability flag worth confirming in a larger DLS run.

#### Thermal stability (DSF)

Thermal unfolding was monitored by intrinsic-fluorescence DSF (dBCM analysis), with trastuzumab again included as a benchmark (Table 4). Trastuzumab’s thermal profile in this assay (Tonset 62.81 °C, Tm1 70.15 °C, Tm2 80.77 °C) is the expected signature of a stable clinical IgG1 and provides a calibration anchor for interpreting the test molecules.

**Table 4.** DSF transition temperatures. Tonset is the temperature at which unfolding begins; Tm1, Tm2 are successive thermal transitions, typically attributed to CH2, CH3/Fab, and Fab core unfolding events in human IgG1.

| Antibody | Tonset (°C) | Tm1 (°C) | Tm2 (°C) |
| --- | --- | --- | --- |
| Converge | 59.54 | 68.34 | 80.96 |
| Cetuximab | 60.89 | 69.49 | 82.72 |
| Cradle | 61.62 | 68.86 | 77.48 |
| Trastuzumab<br>(control) | 62.81 | 70.15 | 80.77 |

All three test antibodies sit within ∼3 °C of trastuzumab’s Tonset and within ∼2 °C of trastuzumab’s Tm1, indicating that none of the molecules present a meaningful thermal-stability liability. The Converge variant’s Tonset (59.54 °C) and Tm1 (68.34 °C) are marginally lower than cetuximab’s (60.89 °C and 69.49 °C, respectively), a ∼1.2 °C decrease consistent with the modest destabilizing effect that can accompany alanine substitutions. Importantly, Converge’s Tm2 (80.96 °C) is essentially equivalent to cetuximab’s (82.72 °C) and to trastuzumab’s reference (80.77 °C), indicating that the Fab domain core remains well folded.

#### Polyspecificity (PSR ELISA)

Polyspecific reactivity is a leading non-target liability for therapeutic antibodies, predicting poor pharmacokinetics, off-target binding, and clearance issues in vivo. We profiled each antibody on a three-substrate PSR ELISA panel (BVP, dsDNA, and insulin) with five reference antibodies that span the polyspecificity scale (Table 5). The control rationale is as follows:

**Table 5.** PSR ELISA results for the three test antibodies and the calibration controls. MFI, median fluorescence intensity at 565 nm; S:B, sample-to-blank ratio; Score, PSR Score (% of Gantenerumab S:B). Lower values indicate lower polyspecificity.

|  | BVP |  |  | dsDNA |  |  | Insulin |  |  |
| --- | --- | --- | --- | --- | --- | --- | --- | --- | --- |
| Antibody | MFI | S:B | Score | MFI | S:B | Score | MFI | S:B | Score |
| Converge | 316,386 | 1.56 | 0.58 | 349,826 | 1.74 | 0.65 | 329,054 | 1.62 | 0.58 |
| Cetuximab | 332,170 | 1.64 | 0.60 | 345,930 | 1.72 | 0.64 | 305,104 | 1.51 | 0.54 |
| Cradle | 534,550 | 2.64 | 0.97 | 638,048 | 3.17 | 1.18 | 472,914 | 2.33 | 0.83 |
| Infliximab (negative PSR) | 313,763 | 1.55 | 0.57 | 414,631 | 2.06 | 0.77 | 298,462 | 1.47 | 0.53 |
| Bevacizumab (low PSR) | 2,533,650 | 12.53 | 4.61 | 3,436,592 | 17.09 | 6.36 | 3,159,544 | 15.59 | 5.57 |
| Bococizumab (high PSR) | 46,525,604 | 230.09 | 84.57 | 54,101,098 | 269.1 | 100.1 | 52,883,862 | 260.8 | 93.19 |
| Gantenerumab (high PSR; ref.) | 55,013,486 | 272.06 | 100 | 54,009,894 | 268.6 | 100 | 56,746,418 | 279.9 | 100 |

- **Infliximab (negative)** — a clinically successful IgG1 with low intrinsic polyspecificity, defining the assay floor.
- **Bevacizumab (medium)** — a clinically approved IgG1 with documented moderate cross-reactivity in PSR ELISA; provides a midpoint anchor.
- **Bococizumab (high)** — a clinical-stage anti-PCSK9 antibody whose program was discontinued in part due to immunogenicity, and which has known PSR liability; defines a “high but not extreme” reference.
- **Gantenerumab (high)** — used as the 100% normalization reference for the PSR Score (PSR Score = 100 × Sample S:B / Gantenerumab S:B).
- **0 nM blank** — background well, used to compute the sample-to-blank ratio (S:B = Sample MFI / 0 nM MFI).

Together, these controls establish a calibration ladder so a sample’s PSR Score is interpretable on a known clinical scale rather than as a raw fluorescence value.

Both Converge and cetuximab are essentially indistinguishable from the negative-control antibody (Infliximab) across all three PSR substrates, with PSR Scores between 0.5 and 0.7 — i.e. below 1% of the Gantenerumab reference and well below the Bevacizumab “medium” anchor (>4.6%). This places both antibodies firmly in the “clean” developability regime for non-specific binding. Cradle shows a consistent ∼1.7-fold elevation across all three substrates relative to Converge and cetuximab; while still well below the medium-control anchor, the systematic elevation across orthogonal substrates is a directional signal worth tracking.

### Summary of developability findings

In aggregate, the Converge variant retains cetuximab’s overall developability. Specifically: (i) DLS hydrodynamic radius and %PD are notably tighter for Converge than for cetuximab, suggesting a more monodisperse, monomeric solution behavior; (ii) HIC retention is modestly elevated relative to cetuximab but well within the developable hydrophobicity envelope; (iii) DSF unfolding transitions track cetuximab within ∼1–2 °C, with no concerning destabilization; and (iv) PSR polyspecificity is statistically indistinguishable from cetuximab and from the clinical negative-control antibody. Across all four assays, no developability red flags were observed for the Converge antibody.

## Discussion

Two findings from this work are worth highlighting. First, ConvergeAB™ — operating in a fully zero-shot mode, with no campaign-specific training and no iteration over experimental data — produced a single lead that improves on a 20-year-old clinical-stage anti-EGFR antibody by ∼2.1× in equilibrium affinity, while also outperforming a contemporary, public-competition-winning design by ∼4.4×. The improvement is achieved with only six conservative substitutions and a global Cα RMSD of 0.15 Å, indicating that the design platform identified a productive minimum on the affinity landscape that is closely adjacent to the cetuximab paratope rather than in some distant sequence space. This is the regime where biobetter design is most valuable: meaningful clinical-grade improvements without abandoning the validated mechanism of action.

Second, the affinity gain comes without a developability cost. Across HIC, DLS, DSF, and PSR ELISA — four orthogonal measurements of hydrophobicity, particle size, thermal stability, and non-specific binding — the Converge variant matches or improves on cetuximab. This is not guaranteed when introducing point mutations into a clinical antibody (alanine substitutions in particular often affect stability), and the fact that the platform’s filtering predictors flagged a candidate that survives all four assays in the lab supports the multi-criterion approach embedded in ConvergeAB™.

The mechanistic picture that emerges is consistent across the data. The four alanine substitutions and two N→D substitutions produce a sub-angstrom local rearrangement at the VL N93A site (0.66 Å Cα shift; 0.5 Å Trp94 displacement) within an otherwise unchanged paratope. Faster k_on_ is consistent with a better pre-organized binding-competent state; slower k_off_ is consistent with improved electrostatic and hydrogen-bond complementarity at the interface. The lower DLS radius and %PD are likewise consistent with a more compact, less conformationally heterogeneous Fv. Together, these observations support a paratope-preorganization model rather than a wholesale binding-mode change.

Several limitations bear acknowledgment. Cell-based potency, ADCC, and in vivo efficacy were not measured here; while improved K_D_ is a strong predictor of these properties for a sterically blocking anti-EGFR antibody, those experiments are an essential next step. Long-term stability (forced-degradation, freeze-thaw, and accelerated stability under formulation conditions) was also not assessed. Finally, the Cradle comparator is a single public sequence and a single competition entry; it should be interpreted as one informative example of contemporary computational design rather than a comprehensive benchmark.

## Conclusion

We demonstrate that ConvergeAB™ can be applied in a fully zero-shot configuration to generate a biobetter antibody with meaningfully improved properties over a clinically established therapeutic. Starting from cetuximab, the platform produced candidates that translated from in-silico predictions to experimentally validated gains in binding kinetics, ultimately yielding an antibody with substantially enhanced affinity to EGFR, and a developability profile that matches or exceeds the parental molecule. These results highlight the ability of ConvergeAB™ to efficiently navigate the antibody design space, and its potential to accelerate the development of next-generation biologics with improved therapeutic performance.

## Methods

### Antibody design (ConvergeAB™)

Cetuximab VH and VL sequences (the affinity-maturation seed) were paired with the recombinant human EGFR extracellular-domain sequence (the target) and submitted to ConvergeAB™ in its default, untrained zero-shot configuration. The platform introduces sequence diversity into the seed antibody using a proprietary protein language model, generating ∼10^5^ target-aware candidates. Candidates are subsequently filtered through a series of explicit, orthogonal predictors that evaluate predicted binding affinity, antibody–target structural compatibility (docking), thermal stability, and solubility. The 10 top-ranked candidates were forwarded to wet-lab validation. Additional platform details are described at https://converge-bio.com/solutions/converge-ab.

### Antibody expression and purification

All three antibodies (Converge, cetuximab, Cradle) and all reference IgG1 controls (trastuzumab, infliximab, bevacizumab, bococizumab, gantenerumab) were produced as full-length human IgG1 with a K214R light-chain substitution to match the clinical form of cetuximab. Production, purification, and quality control were performed at Biointron. Final samples were characterized for identity and purity prior to functional and biophysical assays.

### Surface plasmon resonance (SPR)

SPR experiments were conducted in a multi-cycle kinetics format using recombinant human EGFR-His as the analyte. Each antibody was captured on the chip surface and probed with EGFR at six successive concentrations spanning ∼0.2 to 6.25 nM (0.195, 0.391, 0.781, 1.5625, 3.125, 6.25 nM; an additional 12.5 nM concentration was included in one of the two repeats). Two independent SPR repeats (#1 and #2) were performed under identical conditions, with all three antibodies measured side-by-side in each repeat to enable direct comparison. Kinetic parameters (kon, koff, KD) were extracted by global 1:1 Langmuir fitting of the association and dissociation phases across all concentrations. Per-concentration kobs estimates were combined with dissociation-phase koff estimates and fit linearly (kobs = kon × [EGFR] + koff) to recover kon; KD was computed as koff / kon. Custom analysis scripts are provided with the source data.

### Hydrophobic interaction chromatography (HIC)

HIC-HPLC was performed on a Shimadzu LC-2050C 3D system using a Thermo MabPac HIC-Butyl column (5 µm, 5 × 100 mm) at 25 °C. Mobile phase A was 1.5 M (NH4)2SO4 + 50 mM K2HPO4 (pH 7.0); mobile phase B was 50 mM K2HPO4 + 25% IPA (pH 7.0). Samples were diluted in PBS prior to injection; injection mass was 30 µg. After two minutes of equilibration in 100% A, a linear 0–80% B gradient was run over 30 min, followed by a 0.5 min ramp to 100% B held for 4 min, before re-equilibration in A. Detection was at 280 nm and 214 nm. Trastuzumab (BioIntron, lot B7432) was included as a reference standard for relative-retention-time normalization (RRT = RTsample / RTtrastuzumab).

### Dynamic light scattering (DLS)

DLS was measured on a Wyatt Dynamics Plate Reader (serial 1039-WPR3, Generation 3; control firmware 4.0.1.6) using 384-well format, with a 815.9 nm laser and a 150° detector angle, at 25 °C. Samples were measured in water; viscosity 1.019 cP, refractive index 1.333. Each test antibody was measured in two replicate wells. Cumulant-derived hydrodynamic radius and polydispersity were calculated using Wyatt-recommended cumulant analysis; regularization analysis was performed using the DYNALS algorithm. Peak radius cutoffs were 0.5–10000 nm; correlation-function cutoffs were 1.5–60000 µs.

### Differential scanning fluorimetry (DSF)

Thermal unfolding was monitored by intrinsic Trp fluorescence. dBCM (derivative of the barycentric mean of the emission spectrum) was plotted vs. temperature, and unfolding transitions (Tonset, Tm1, Tm2) were extracted from the resulting trace. Trastuzumab was included as a thermal-stability benchmark control.

### Polyspecificity reagent (PSR) ELISA

PSR ELISA was performed on three orthogonal substrates: bovine viral particle (BVP), double-stranded DNA (dsDNA), and insulin. Substrates were coated overnight at 4 °C, followed by a wash, blocking (1 h, RT), and three further washes. Each test antibody was applied as primary antibody at 100 nM (1 h, RT) followed by six washes. Detection was performed with goat anti-human IgG-Fc HRP (Sigma A0170, 1 h, RT), six washes, and Amplex Red substrate read at 565 nm fluorescence on a Molecular Devices SpectraMax i3X plate reader. Five reference IgG1s were included as calibration controls: Infliximab (negative), Bevacizumab (medium), Bococizumab (high), and Gantenerumab (high; used as the 100% normalization reference). PSR Score was computed as 100 × Sample S:B / Gantenerumab S:B, where S:B = Sample MFI / 0 nM MFI.

## Author contributions

I.W. conceptualized the study, prompted the ConvergeAB™ platform, designed the experiments, analyzed the results, and wrote the manuscript.

## Data availability

Source data are available upon reasonable request.

## Acknowledgements

We thank Professor Angel Progador for critical reading of the manuscript. We also thank Biointron for their support with laboratory work.

## Conflict of Interest

The author, Iddo Weiner, serves as Chief Scientific Officer (CSO) of Converge Bio Ltd.

